# MmCMS: Mouse models’ Consensus Molecular Subtypes of colorectal cancer

**DOI:** 10.1101/2022.06.17.496539

**Authors:** Raheleh Amirkhah, Kathryn Gilroy, Sudhir B Malla, Tamsin RM Lannagan, Ryan M Byrne, Natalie C Fisher, Shania M Corry, Hojjat Naderi-Meshkin, Baharak Ahmaderaghi, Richard Murray, Megan Mills, Andrew D. Campbell, ACRCelerate consortium, Antoni Berenguer Llergo, Rebeca Sanz-Pamplona, Alberto Villanueva, Eduard Batlle, Ramon Salazar, Mark Lawler, Owen J. Sansom, Philip D. Dunne

## Abstract

**BACKGROUND:** Colorectal cancer (CRC) primary tumours are molecularly classified into four consensus molecular subtypes (CMS1-4). Genetically engineered mouse models aim to faithfully mimic the complexity of human cancers and, when appropriately aligned, represent ideal pre-clinical systems to test new drug treatments. Despite its importance, dual-species classification has been limited by the lack of a reliable approach. Here we utilise, develop and test a set of options for human-to-mouse CMS classifications of CRC tissue.

**METHODS:** Using transcriptional data from established collections of CRC tumours, including human (TCGA cohort; n=577) and mouse (n=57 across n=8 genotypes) tumours with combinations of random forest and nearest template prediction algorithms, alongside gene ontology collections, we comprehensively assess the performance of a suite of new dual-species classifiers.

**RESULTS:** We developed three approaches: MmCMS-A; a gene-level classifier, MmCMS-B; an ontology-level approach and MmCMS-C; a combined pathway system encompassing multiple biological and histological signalling cascades. Although all options could identify tumours associated with stromal-rich CMS4-like biology, MmCMS-A was unable to accurately classify the biology underpinning epithelial-like subtypes (CMS2/3) in mouse tumours.

**CONCLUSIONS:** When applying human-based transcriptional classifiers to mouse tumour data, a pathway-level classifier, rather than an individual gene-level system, is optimal. Our R package with three options helps researchers select suitable mouse models of human CRC subtype for their experimental testing.

## INTRODUCTION

Colorectal cancer (CRC) primary tumours can be molecularly classified into four consensus molecular subtype (CMS1-4) (1). According to this classification, CMS1 (14% of patients) is enriched for tumours with microsatellite instability (MSI) and immune activation. CMS2 (37% of patients) epithelial-rich tumours represent the canonical subtype and are associated with activation of the WNT/MYC pathways and chromosome instability. CMS3 (13% of patients) tumours display signalling indicative of increased metabolic activity and *KRAS*-mutations. Finally, CMS4 (23% of patients) tumours display stromal-rich and mesenchymal features, alongside activation of TGF-β and VEGFR pathways (1). While CMS classification provides valuable prognostic information, its ability to identify subtype-specific responses to therapies remains an area of active research, with several reverse-translation studies using human pre-clinical models, such as cell lines, organoids and patient-derived xenografts (PDX) (2,3). While CMS classification in these models is possible, the reliance of CMS classification on gene expression signals from tumour microenvironment compartments can undermine attempts to identify the mesenchymal subtype of CRC (CMS4) in cell lines, patient-derived organoids and PDXs (4,5). To address this, Eide and colleagues developed a CMS classifier specifically designed for human pre-clinical models, named CMScaller, which used a filtered set of cancer cell-intrinsic, subtype-enriched gene expression markers, giving a surrogate measurement of alignment with CMS subtypes in *in vitro* and *in vivo* models (6).

Although translation of human CMS subtypes to human-based pre-clinical models has been addressed, there remains a need to develop and test a classifier that can be used with mouse-based tumour data from genetically engineered mouse models (GEMMs). GEMMs, alongside the armament of human pre-clinical models, represent the most appropriate models to mimic the complexity of the human CRC biology. GEMMs in particular provide an ideal system to improve pre-clinical drug testing within a native immunocompetent host (7,8). Identifying murine models that recapitulate each CRC subtype features can therefore de-risk clinical translation of therapeutics, while also providing an excellent opportunity to improve our understanding of the nuanced and complex interactions between cancer epithelial cells and their microenvironment. Currently, there is no reliable and standardised approach for CMS classification using data from mouse tissues. In the absence of such a system, users have relied on converting the human CMS template to mouse orthologues, followed by sample classification using the nearest template prediction (NTP) method (as with the CMScaller), or conversely converting mouse genes to human orthologues and applying the random forest (RF) method used in the original CMSclassifier algorithm (1,6). Both approaches rely on overlapping nomenclature for individual genes; as mouse genes with different names to the ones in the human classifier template will be ignored/removed during CMS assignment, or vice versa. In addition, both systems are also fully reliant on the assumption that genes within the classifier will perform the same biological function in both mouse and human tumours and ignore interspecies variability. Recent studies have shown that pathway-based classifications are more robust as they are composed of tens to hundreds of co-ordinately expressed genes, and therefore are protected to some degree from the loss of individual genes or variations in functions, both of which are known to undermine gene-level classifiers (9,10). As such, pathway-based approaches consider the collective impact of genes on pathway-level activity rather than being influenced by a single differentially expressed gene. Furthermore, broad biological knowledge-based approaches have previously been shown to be less influenced by non-biological factors (11,12).

To improve on the current state-of-the-art approach of classifying GEMM tumours, we developed three options for CMS classification in mouse tissue. The first, hereafter named as MmCMS-A, uses mouse orthologues of the human CMS gene template from CMScaller (6), thus it has sole emphasis on individual genes. Given the benefits of pathway-level approaches for classification, over gene-level, we proposed two further options (MmCMS-B and MmCMS-C) that use biological knowledge-based information from either gene ontology (MmCMS-B) or a compendium of signatures from biological signalling collections and microenvironment populations (MmCMS-C). Most importantly, to ensure the field can utilise these mouse CMS classification approaches, we developed an R package, namely MmCMS, which provides a publicly-available tool to classify samples according to all three options, enabling users to assess the alignment of GEMM tumours to human CMS subtype.

## METHODS

### Human CRC cohort

The processed TCGA COREAD RNA-Seq dataset (n = 577) was downloaded directly from the Guinney et al., *CMSclassifier* study via Synapse (ID: syn2023932), where it has been described previously (1). Gene symbols and entrez IDs were matched using *org*.*Hs*.*eg*.*db* R package (v3.8.2) thereafter CMS classification was performed via ‘random forest’ (RF) method using *CMSclassifier* R package (version 1.0.0).

Biological Process subset of Gene Ontology (GO) gene sets were extracted from the Molecular Signature Database (MSigDB) using *msigdbr* R package (v7.0.1). Subsequently, ontology scores were generated for the TCGA dataset using single sample GSEA (ssGSEA) method from *GSVA* R package (v1.26.0). To determine the CMS-specific GO terms, these ssGSEA scores were averaged for each gene set across samples within each CMS subtype and scaled to Z-scores where the GO with ssGSEA scaled scores above 0 in a CMS, but below 0 on the others, were selected as the enriched GO term for that CMS. The CMS-specific GO BP gene sets for mouse species were then extracted from the msigdbr R package and used to develop an ontology-based CMS classification for mouse.

### Mouse models

All animal experiments were performed in accordance with a UK Home Office licence (Project License 70/8646), and were subject to review by the animal welfare and ethical review board of the University of Glasgow. Mice of both sexes were induced with a single injection of 2 mg tamoxifen (Sigma-Aldrich, T5648) by intraperitoneal injection at an age of 6 to 12 weeks, all experiments were performed on a C57BL/6 background. Mice were sampled at clinical endpoint, which was defined as weight loss and/or hunching and/or cachexia.

### Mouse RNA sequencing and analysis

RNA was isolated using either an RNeasy mini kit (Qiagen) or TRIzol reagent (Thermo Fisher Scientific). RNA concentrations were determined using a NanoDrop 200c spectrophotometer (ThermoScientific), and quality was assessed using an Agilent 220 Tapestation using RNA screentape. RNA sequencing was performed using an Illumina TruSeq RNA sample prep kit, then run on an Illumina NextSeq using the High Output 75 cycles kit (2 x 36 cycles, paired end reads, single index). Raw sequence quality was assessed using the FastQC algorithm version 0.11.8. Sequences were trimmed to remove adaptor sequences and low-quality base calls, defined as those with a Phred score of <20, using the Trim Galore tool version 0.6.4. The trimmed sequences were aligned to the mouse genome build GRCm38.98 using HISAT2 version 2.1.0, then raw counts per gene were determined using FeatureCounts version 1.6.4. Raw read counts of the small cohort (n=18) which is publicly available at ArrayExpress: E-MTAB-6363 were normalized using vst function in DESeq2 R package (v1.32.0). The models where the batch they were sequenced in was deeply confounded by genotype were removed and data from 51 GEMMs remained. ComBat_seq function in sva R package (v3.40.0) was used to correct read counts for batch, thereafter vst function in DESeq2 same as before was used to normalize the data.

### Databases

CMS curated gene sets signatures (n=79) were obtained from Synapse (ID: syn2321865). Cancer hallmarks (n=50) and Gene Ontology (GO) Biological Process gene set (C5 BP) was extracted from MSigDB using ‘msigdbr’ R package (v7.4.1).

Ten signatures to estimate the proportion of the eight immune (NK cells, Cytotoxic lymphocyte, T cells, CD8 T cells, B lineage, Monocytic lineage, Neutrophils, Myeloid dendritic cells) and two stromal (Fibroblasts and Endothelial) cell populations in each human sample across CMS subtypes were obtained using the MCPcounter R package (v1.2.0); the mouse version of signatures were retrieved from the mMCPcounter R package (v0.1.0). Immune-related genes for human and mouse were downloaded from the NanoString panel (https://canopybiosciences.com/product/immunology/).

### Statistical analysis

All the statistical analyses were performed in R (v4.1.2) using the *stats* R package, including *cor()* function with method = ‘pearson’ for Pearson’s correlation. The Student *t-test* method embedded in the *geom_signif()* function of *ggsignif* package (v0.6.3) was used to do statistical analysis in violin plots. Boxplots were generated using *ggplot2* (v3.3.5) R package. The *ComplexHeatmap* (v2.8.0) and *circlize* (v0.4.13) packages were used to display heatmaps. We used *glmnet* (v4.1-3) R package to do LASSO regression model analysis. The λ or tuning parameter in the LASSO model was selected through the 10-fold cross-validation. Transcriptome-based stromal and immune scores were generated using the *MCPcounter*.*estimate()* function in *MCPcounter* R package (v1.2.0). Single sample gene set enrichment analysis (ssGSEA) was performed using an R package called *GSVA* (v1.40.1). Alluvial plot to display concordance result was drawn using *riverplot* (v0.10). The Nearest Template Prediction (NTP) algorithm, with cosine correlation distances, was employed to predict the proximity of each GEM model’s expression profile to the four CMS subtypes, using each of the three templates individually (A, B and C), with an FDR <0.05 used as a cut off for statistical significance.

## Results

### Development and testing of CMS classifier templates for use in mouse tumours

To confirm the concordance between the CMScaller/pre-clinical CMS classifier (NTP method) and the CMSclassifier/original CMS classifier (RF method) in human data, we applied CMScaller on COREAD TCGA RNA-seq transcriptional data (n=577) retrieved from the original CMS article (1). After removing samples that were unclassified by either RF or CMScaller, we found 91.19% (321/352) concordance between RF and NTP calls (Supplementary Fig. 1a). PCA analysis on the whole transcriptome of these 352 samples demonstrated that samples that gave conflicting calls (indicated as swapped in Supplementary Fig. 1a) between RF and CMScaller were in the boundary of CMS subtypes assigned by the RF method (Supplementary Fig. 1a). To confirm that the discrepancies in classification call were confined to samples with lower CMS probability scores, when we set a more stringent CMS classification probability cut off (>0.8) for the RF method, the classifications for the two methods increased to 100% concordance, n=93 (Supplementary Excel File 1, Sheet 1), demonstrating that CMScaller provides excellent CMS classification concordance for samples that display the strongest CMS transcriptional traits, as indicated by high subtype RF classification scores.

While these data confirm the suitability of using either the RF CMSclassifier or NTP CMScaller methods for CMS classification of human tumour data, to assess the performance of these methods on mouse tumour model classification, we next assembled transcriptional data from two independent GEMM tumour cohorts (Table 1). Tamoxifen-regulated Cre-*loxP* system was used to generate all models and introduced via an intraperitoneal injection. The small cohort has been previously described by Jackstadt et al. and composed of 18 intestinal primary tumours across 4 genotypes that represent both the serrated (KPN: *Kras*^G12D/+^ *Trp53*^fl/fl^ *Notch1*^Tg/+^; KP: *Kras*^G12D/+^ *Trp53*^fl/fl^) and tubular (APN: *Apc*^fl/+^ *Trp53*^fl/fl^ *Notch1*^Tg/+^; AP: *Apc*^fl/+^ *Trp53*^fl/fl^) tumour histologies (13). The large independent cohort (n=39) contained a set of independent KP and KPN tumours alongside 4 additional genotypes including *Apc*^fl/+^ (A); *Apc*^fl/+^ *Kras*^G12/+^ (AK); *Braf*^V600E/+^ *Trp53*^fl/fl^ (BP) and *Braf*^V600E/+^ *Trp53*^fl/fl^ Notch1^Tg/+^ (BPN). Median latency age of A, AK AP, APN, KP, KPN, BP and BPN models is 215, 67, 185, 161, 171, 184, 190 and 174 days respectively, developing small intestine (SI) tumours primarily, with the exception of seven mice (AK=5, A=2) which formed tumours in colon. For more characterisation of the samples see Supplementary Excel File 2.

**Table 1.** Summary of mouse models used in this study

|  | Model name | Genotype of mouse model |
| --- | --- | --- |
| **Small cohort (n=18)** | **AP** | villinCre<sup>ER</sup> Apc<sup>fl/+</sup> Trp53<sup>fl/fl</sup> (n=3) |
|  | **APN** | villinCre<sup>ER</sup> Apc<sup>fl/+</sup> Trp53<sup>fl/fl</sup> Notch1<sup>Tg/+</sup> (n=3) |
|  | **KP** | villinCre<sup>ER</sup> Kras<sup>G12D/+</sup> Trp53<sup>fl/fl</sup> (n=3) |
|  | **KPN** | villinCre<sup>ER</sup> Kras<sup>G12D/+</sup> Trp53<sup>fl/fl</sup> Notch1<sup>Tg/+</sup> (n=9) |
| **Large cohort (n=39)** | **A** | villinCre<sup>ER</sup> Apc<sup>fl/+</sup> (n=6) |
|  | **AK** | villinCre<sup>ER</sup> Apc<sup>fl/+</sup> Kras<sup>G12D/+</sup> (n=6) |
|  | **BP** | villinCre<sup>ER</sup> Braf<sup>V600E/+</sup> Trp53<sup>fl/fl</sup> (n=4) |
|  | **BPN** | villinCre<sup>ER</sup> Braf<sup>V600E/+</sup> Trp53<sup>fl/fl</sup> Notch1<sup>Tg/+</sup> (n=7) |
|  | **KP** | villinCre<sup>ER</sup> Kras<sup>G12D/+</sup> Trp53<sup>fl/fl</sup> (n=6) |
|  | **KPN** | villinCre<sup>ER</sup> Kras<sup>G12D/+</sup> Trp53<sup>fl/fl</sup> Notch1<sup>Tg/+</sup> (n=10) |

The RF method in the CMSclassifier package was designed for human samples and uses 273 genes to assign CMS subtypes. To enable the use of this method with mouse data, we converted the entire mouse gene matrix to human orthologues using biomaRt (14). During the conversion of the mouse matrix, 16 genes of the 273 gene used to predict CMS calls in human were mismatched in both cohorts (Supplementary Table S1). Applying the RF method to our n=18 and n=39 mouse model matrices produced 56% unknown samples in both datasets (Supplementary Fig. 1b, c). Of note, to test the functionality of CMScaller in the same mouse cohorts, we next converted the human CMScaller template genes (n= 529; CMS1=126, CMS2=82, CMS3=84, CMS4=237) to mouse orthologues (n=533; CMS1=128, CMS2=80, CMS3=90, CMS4=235), which as anticipated resulted in a small number of dropouts (n=26 missing genes, Supplementary Table S2) due to lack of recognised orthologues, though overall the number of genes in mouse CMS template increased due to the existence of multiple mapping mouse genes for the individual human genes (Fig. 1a, Supplementary Excel File 3). Using this CMScaller method in our mouse data, we found fewer unknown samples, 17% and 36%, respectively (Supplementary Fig. 1b, c) and therefore selected this NTP-based approach as our initial dual-species classifier, termed MmCMS-A.

**Fig. 1:**
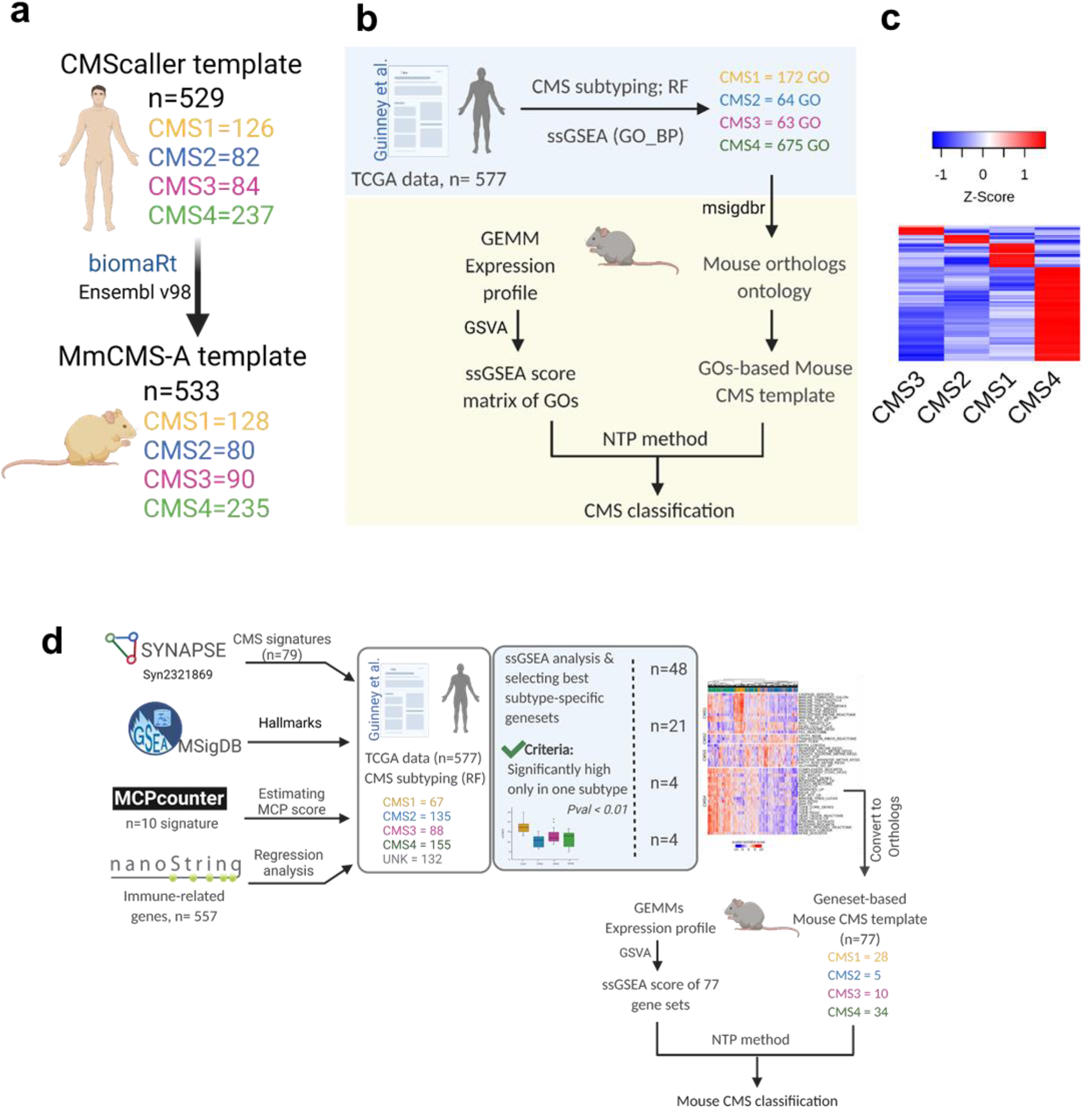
Three different approaches for mouse CMS subtyping. **a** The schematic shows the approach of converting the human CMS template to mouse orthologues (MmCMS-A option). **b** Schematic of developing a gene ontology-based classifier to call CMS subtypes in mouse tissues. **c** Heatmap of ssGSEA for selected GO-BP terms based on z-score > 0 in a CMS subtype and z-score < 0 for other subtypes in human data. **d** Schematic of developing MmCMS-C classifier based on four biologically-informed signature collections (Figure 2a heatmap shown as an exemplar).

### Identification of CMS-related GO-BP terms in human TCGA data (MmCMS-B)

To complement the gene-level approach in MmCMS-A, we again utilised the RF classifications used in the original CMS classifier development within the human TCGA COREAD data (n=577) to identify the gene ontology (GO)-biological process (BP) that are most significantly associated with individual human CMS classes, using gene set enrichment analysis at single sample level (ssGSEA) to derive the enrichment score across all samples. For each gene set (GO-BP), the mean enrichment score was calculated for each human CMS subtype, with scaled Z-score > 0 in one CMS subtype but < 0 in the other three CMS subtypes being selected as distinct features for each particular CMS subtype. This identified n=172, n=64, n=63 and n=675 specific GO terms associated with CMS1, CMS2, CMS3 and CMS4 respectively (Fig. 1b, blue background; Fig. 1c). To test if these CMS class-specific GO-terms represent surrogate markers for human samples called with high probability using RF, we again used CMS classifications from the stringent RF threshold (probability cut-off > 0.8) as before and compared them to CMS classifications using this new NTP ontology-based method, where we observe 95% concordance with the RF-based calls (Supplementary Excel File 1, Sheet 2). In line with the generation of MmCMS-A, the mouse-equivalent GO-terms of these human gene ontologies were identified using the ‘msigdb’ R package and used as the MmCMS-B template for CMS classification in mouse data using NTP method (Fig. 1b, yellow background).

### Development of mouse CMS template (MmCMS-C) based on combining gene sets/ pathways that best characterise each human CMS subtype in a supervised approach

While MmCMS-B is focussed solely on GO-BP signatures, for MmCMS-C we generated a classifier based on four biologically-informed signature collections (Fig. 1d). First, we compiled the n=79 gene sets used to characterise biological signalling in the original CMS study from the Synapse database (DOI: https://doi.org/10.7303/syn2623706). As with MmCMS-B, we refined these 79 signatures into only those with individual CMS class-specific expression (t.test; pvalue < 0.01) and signatures only kept if one subtype was significantly higher when compared to each of the other subtypes in turn, resulting in 48 of the 79 gene sets being used (Fig. 2a; Supplementary Fig. 2a). Next, using the 50 MSigDB hallmark gene sets, we identified 21 with significant expression (t.test; pvalue < 0.01) across CMS groups (Fig. 2b; Supplementary Fig. 2b). In the third step, we used the microenvironment cell population (MCP)-counter signatures, and in line with previous studies, we found cytotoxic lymphocyte and NK cells are significantly enriched in CMS1, whereas fibroblast and endothelial cells are enriched in CMS4, thus 4 signatures from MCPcounter (15) were included (Supplementary Fig. 2c; t.test; pvalue < 0.01). Finally, given the importance of inflammatory lineages in development and classification, we assessed immune-related genes (n= 557; from a NanoString panel) for their associations with each CMS subtype, filtered first using the Least Absolute Shrinkage Selector Operator (LASSO) regression model (Fig. 2c). Based on coefficient > 0, overall 44 immune-related genes (CMS1=14, CMS2= 8, CMS3= 9, CMS4= 13) were found as the best predictors of individual CMS classes. As with Options B, these co-ordinately expressed immune genes for each CMS subtype were then grouped for ssGSEA, and enrichment scores were assessed across subtypes which were significantly enriched (t-test; pvalue <0.01) (Fig. 2c). Overall, this four-step MmCMS-C approach identified 77 CMS class-specific gene sets (CMS1=28; CMS2=5; CMS3=10; CMS4=34). When tested in the same way as MmCMS-A and B, using the NTP method on TCGA data, MmCMS-C was found to have 98% concordance with the RF-based high probability calls, threshold = 0.8 (Supplementary Excel File 1, Sheet 3). To enable mouse classification, the biomaRt (14) and msigdb (16) R packages were used to obtain the mouse version of 48 gene sets and 21 hallmark pathways, respectively, with mouse MCP signatures retrieved from the mouse-specific mMCP-counter package (17). Individual orthologues of immune-related genes were obtained from the mouse NanoString panel (Supplementary Table S3), with 39 mouse genes aligned to the 44 human immune genes identified using regression analysis, and then grouped into signature scores as before.

**Fig. 2:**
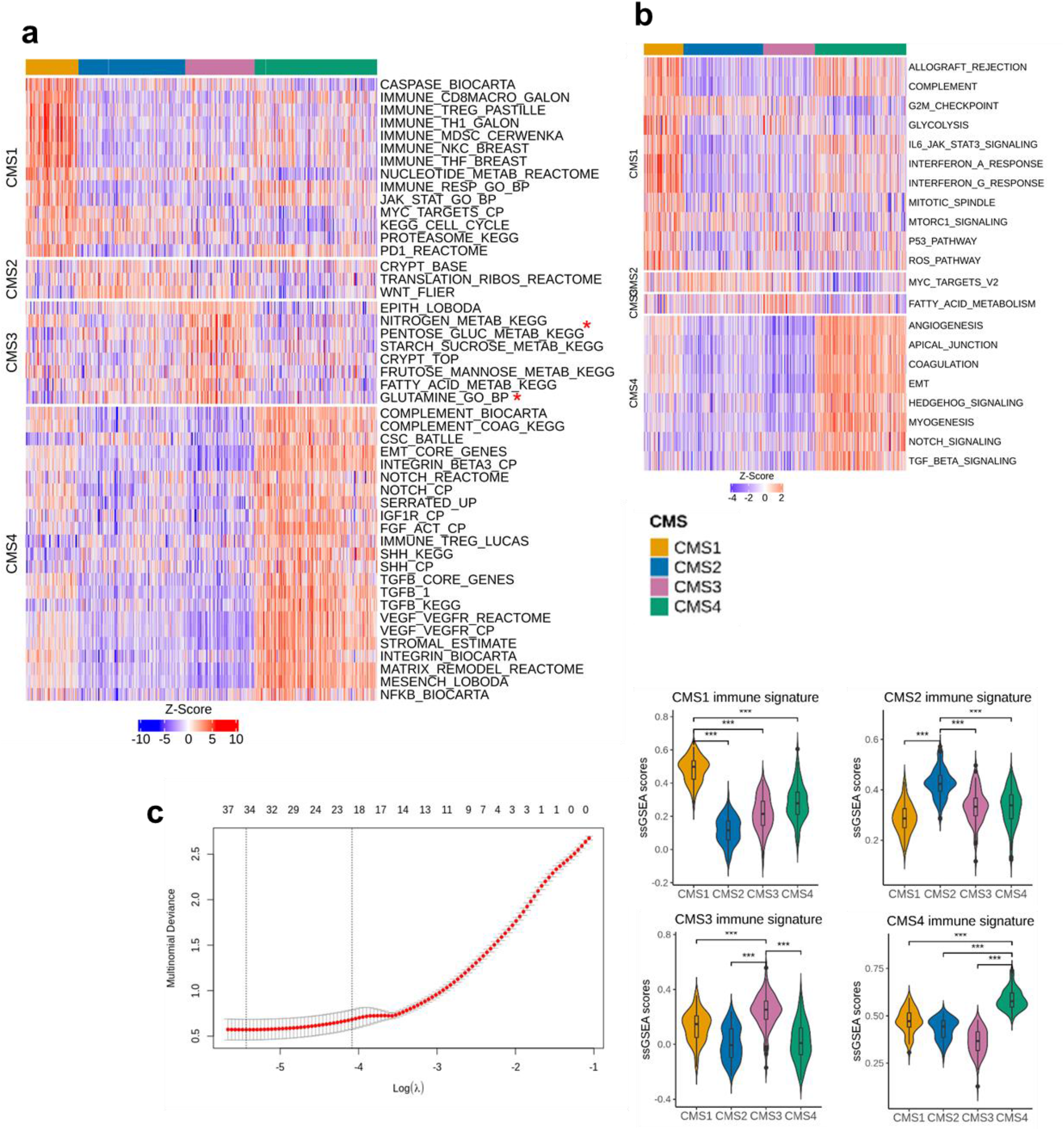
Identification of pathways and biology that are best characteristic of each human CMS subtype for Option C. **a** ssGSEA scores heatmap of 48 CMS-related signatures (CMS1=14, CMS2=3, CMS3=8, CMS4=23), from original the CMS article(1), that are significantly different (pairwise t-test; see Supplementary Fig. 2b) across CMS subtypes in human dataset. Scores are converted to Z scores. * PENTOSE_GLUC_METAB_KEGG and GLUTAMINE_GO_BP are defined with different names in the CMS curated gene sets signatures (n=79) from Synapse (ID: syn2321865) but the genes are the same. **b** ssGSEA scores heatmap of 21 selected hallmark pathways that are significantly different (pairwise t-test; see Supplementary Fig. 2b) across CMS subtypes in the human dataset. Scores are converted to Z scores. **c** Selection of the λ parameter in the LASSO model by 10-fold cross-validation based on minimum criteria. Red dots show the average deviance values for each model with a given λ. The vertical black lines define the optimal values of λ, where the model provides the best fit to the data. A λ value of 0.01682636 (lambda.1se), was chosen. Violin plot and pairwise t-test used to display enrichment of selected immune gene set across CMS subtypes (CMS1 n=14, CMS2= 8, CMS3= 9, CMS4= 13). *** p ≤ 0.001

### CMS classification of GEM models using 3 options

To assess the performance of our 3 options for classifying mouse tumours, two different cohorts of GEMMs as described above were used (Table 1, Fig. 3a). As there is no CMS “ground truth” or reference for mouse tumour data, we utilised tumours from n=18 mouse models across four genotypes (KPN, KP, APN, AP; Table 1), which we have previously shown to correlate with signalling associated with stromal CMS4 tumours (KPN and KP) or epithelial-rich CMS2/3 tumours (AP and APN). PCA on the dataset revealed distinct groups according to genotype (Supplementary Fig. 3a). The NTP-based algorithm was employed to predict CMS classification of GEMM tumours, using each of the three templates individually (MmCMS-A, B and C), with an FDR <0.05 used as a cut off for significant calls. Within the small cohort, both MmCMS-A and MmCMS-C returned 3 unknown calls, however MmCMS-B classified all mouse tumours (Fig. 3b). Although some intra-genotype variation in CMS classifications were identified, indicating heterogeneity within tumours with the same genotype, these findings were all in line with previously published subtype associations for these models (Fig. 3b and Supplementary Table S4).

**Fig. 3:**
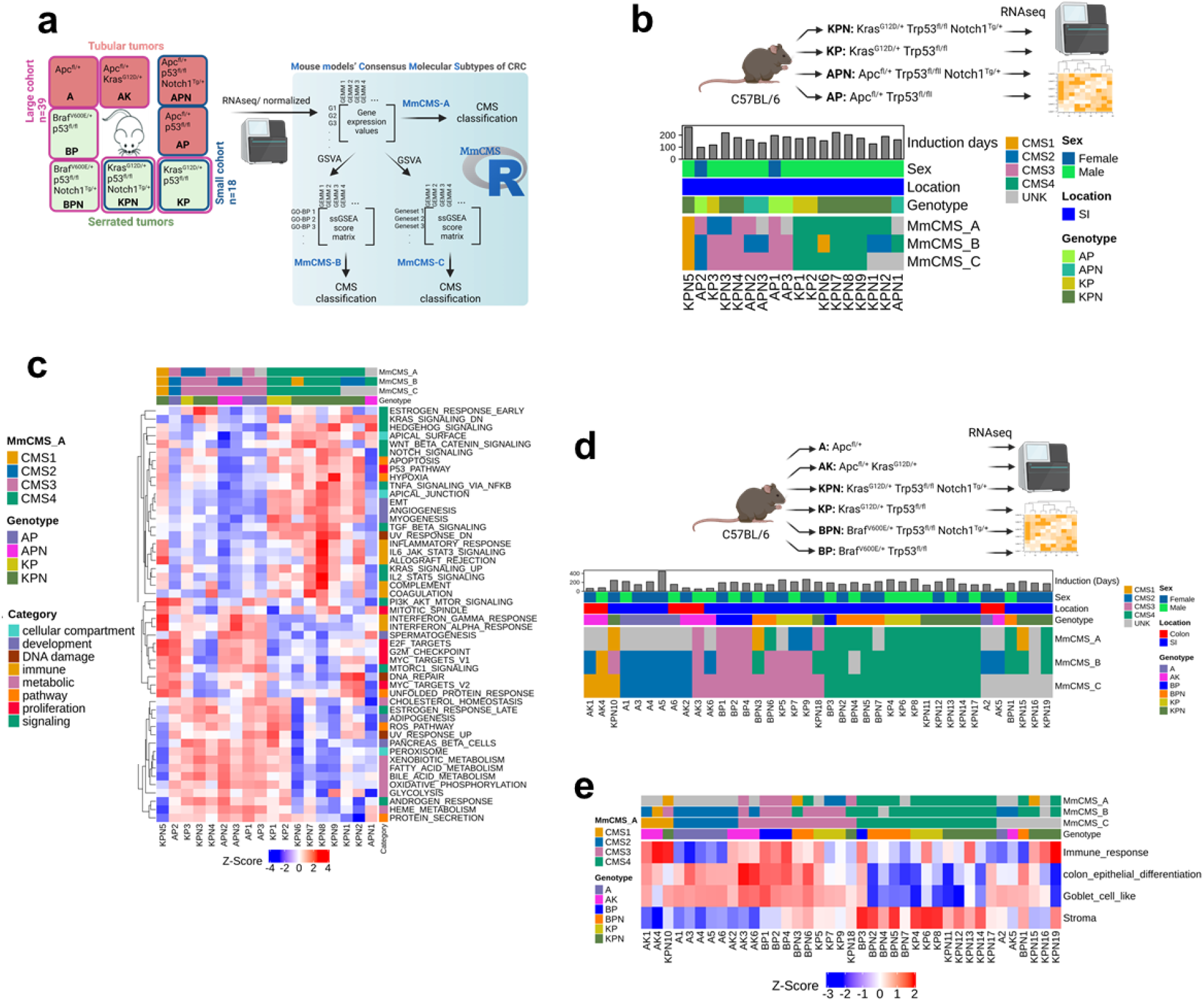
Molecular characterization of GEM models. **a** Left part of schematic shows all the GEMMs used in this study. The border colour indicates the genotypes that are included in each cohort, blue for small cohort and pink for large cohort. The squares with both border colours show the presence of that particular genotypes in both cohorts, but the samples are different. The squares with peach background are representative of the tubular tumour models and green background show the serrated tumour models. Right part of schematic shows the input file for each three options. MmCMS-A option use normalized expression values to call CMS for mouse tissues, but MmCMS-B and -C work on ssgsea score matrix. All the analysis process to convert gene expression values to ssgsea score matrix will be automatically done in the R package. Users just need to provide the package with normalized expression values as input file while genes are row names and samples are in columns. **b** Comparison of CMS classification results using our three options (MmCMS-A, MmCMS-B and MmCMS-C) in small GEMM cohort (n=18). Grey colour indicates unclassified samples. The CMS calls are aligned by genotype, location, sex and duration of tamoxifen induction. **c** Heatmap of Hallmark ssGSEA score across samples from the small cohort of GEMMs (scores are z-score scaled). **d** CMS classification of 39 GEM models using our three options. Grey colour indicates unclassified samples. The CMS calls are aligned by genotype, tumour location, sex and duration of tamoxifen induction. **e** ssGSEA scores heatmap of immune response, colon epithelial differentiation, goblet cell–like, and stroma related gene sets across GEMMs that are aligned by genotype and CMS classification result from the 3 options in the large cohort (n=39).

There was broad consensus across all three options for samples classified as CMS4, indicating how distinct this subtype is compared to the others, however samples classified as CMS3 using MmCMS-C were classified as either CMS2 or unclassified using MmCMS-B and MmCMS-A (Fig 3b). Characterisation of these GEMM tumours, using ssGSEA, shows that as with human tumours, all samples assigned as CMS4, most prominently by MmCMS-C, display the highest levels of enrichment for TGF signalling, EMT, angiogenesis, Notch and Hedgehog signalling. In line with human CMS biology, samples classified as CMS3 using MmCMS-C display high expression of metabolic pathways, such as bile acid, xenobiotic, fatty acid, heme metabolism and glycolysis. Samples classified as CMS2 have high expression of MYC and E2F targets which are well-identified signalling molecules in the CMS2 subtype. One sample was consistently classified as CMS1, which displayed high expression of interferon-gamma response and interferon-alpha response (Fig. 3c and Supplementary Fig. 3b).

Following assessment in this initial cohort of histologically distinct tumours, with tubular/epithelial-rich (AP, APN) and serrated/stroma-rich (KP, KPN) genotypes, we next tested each of the individual classifier options in an independent and more heterogeneous cohort of 39 mouse tumours across 6 genotypes. When applied using the same method as above, MmCMS-A, B and C returned n=14, n=2 and n=6 unknown samples respectively (Fig. 3d). MmCMS-A returned unknown calls for all *Apc*^fl/+^ samples, however when using MmCMS-B and C all samples with *Apc*^fl/+^ genotype were assigned as CMS2, with biological characterisation using GSEA indicating that these samples have enriched signalling hallmarks related to proliferation including G2M checkpoint, E2F targets and MYC targets (Fig. 3d). The only *Apc*^fl/+^ genotype sample (A2) that remained unclassified by MmCMS-C appeared as an outlier when assessed by PCA, as it did not cluster with other *Apc*^fl/+^ samples (Supplementary Fig. 3c).

GSEA reveals that samples classified as CMS2 by MmCMS-A display features inconsistent with human CMS2 tumours, and are more aligned with human CMS3 classification, including high expression of metabolic pathway and low expression of proliferation-related hallmarks, indicating limited ability of the gene-level approach to identifying CMS2 tumours (Fig. 3d, Supplementary Fig. 3b). Furthermore, genes associated with immune response, colon epithelial differentiation, goblet cell–like, and stroma, which represent CMS1, CMS2, CMS3, and CMS4, respectively, were obtained from (18), converted to mouse orthologues using biomart and examined in the GEMMs to determine if they support the CMS calls assigned by the classifiers (Fig. 3e and Supplementary Fig. 3d). Referring to MmCMS-C calls particularly, this analysis reveals a strong association between CMS4 samples and stroma signature. The CMS2 samples in the larger cohort (n=39) are repressed for immune response and stroma signatures but have high enrichment for colon epithelial differentiation as expected as well as goblet cell-like signatures (Fig. 3e). Although all CMS3 samples have universal enrichment for goblet cell-like signatures, some samples with BP, BPN, AK genotype also display elevated immune response and colon epithelial differentiation signatures. Moreover, the result demonstrates high enrichment of only immune response signature for CMS1 samples in the small cohort as expected, however in the larger cohort there is also some level of expression for colon epithelial differentiation and goblet cell-like signatures, although these inconsistencies may be explained due to limited samples classified as CMS1 using any method (Fig. 3e and Supplementary Fig. 3d).

### MmCMS-C is the best option, particularly in calling CMS2-like mouse tissues

To test how well our GEMM classifications align to the biological characteristics associated with human CMS subtypes, we next measured the biological traits of immune-related, metabolic, proliferation and stromal signalling associated with CMS calls in human TCGA data and compared them directly to the CMS classification calls according to each of our three MmCMS options in both independent GEMM cohorts. Mean ssGSEA scores were calculated across samples of each human CMS subtype, using the same TCGA samples used in Figure 1, alongside mean ssGSEA scores for MmCMS-A, B and C predictions in the n=18 and n=39 GEMM cohorts (Fig. 4a-b).

**Fig. 4:**
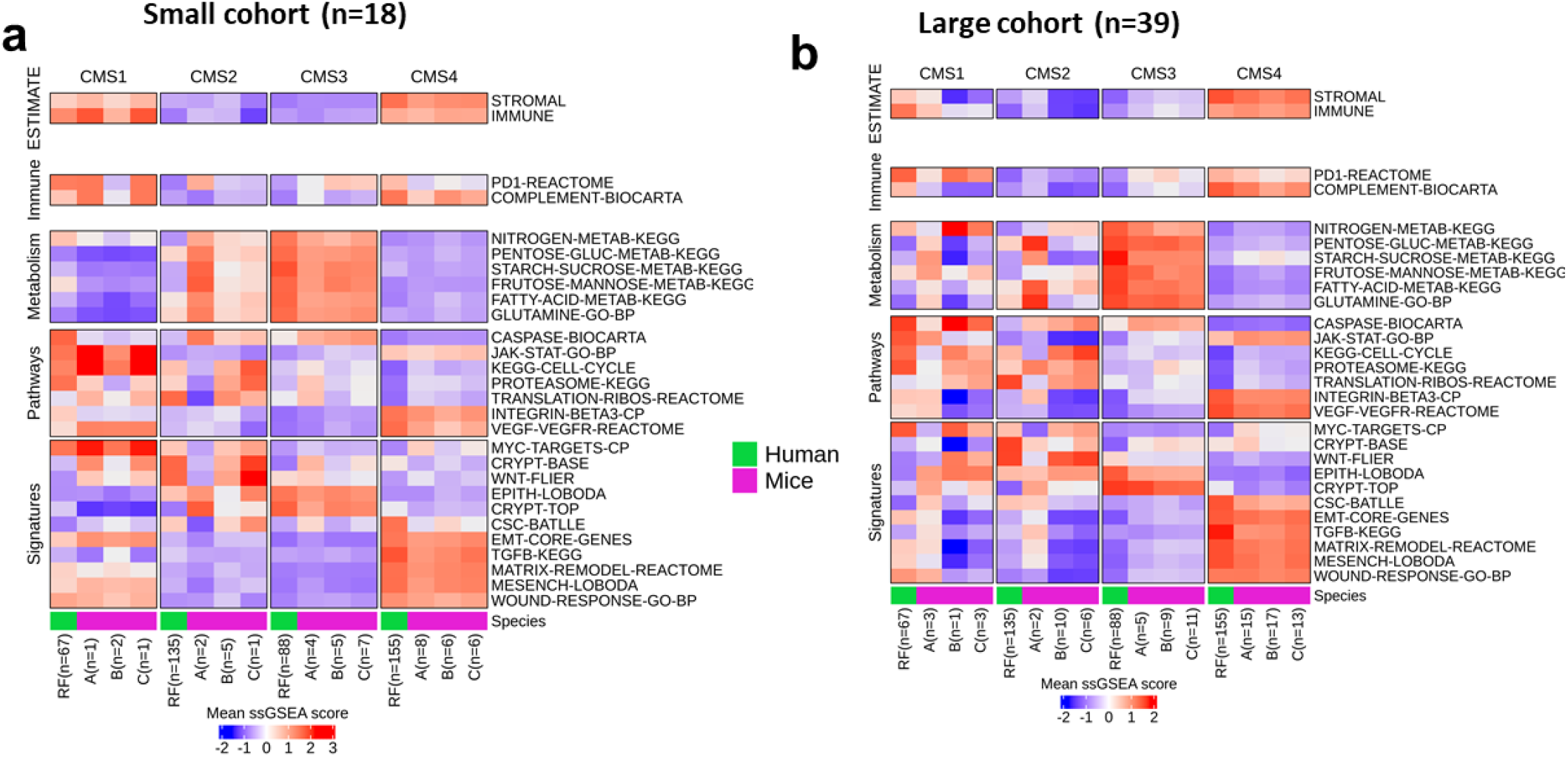
Pathway-based classification is more reliable, particularly in calling CMS2-like mouse tissues. **a** Comparison of mean ssGSEA scores from the biological characteristics associated with human CMS subtypes, applied in mouse CMS calls (n=18 GEMM cohort) using the 3 classification options and human CMS calls. **b** Comparison of mean ssGSEA scores of hallmarks in mouse CMS calls (n=39 GEMM cohort) using the 3 classification options and human CMS calls.

Using the human RF calls as the ground truth, followed by cross-comparison and correlation analysis of samples assigned as CMS2 by all three mouse options, we find strong correlation with MmCMS-B (r=0.79, p=0.0000005) and MmCMS-C (r=0.81, p=0.0000001) and no correlation with MmCMS-A in both cohorts (Fig. 4, Supplementary Table S5). Additionally, we found limited associations for biological traits in human CMS1 with the CMS1 calls for any of our mouse classifier options, again however this may be due to small numbers of CMS1 classifications in mouse tumours. In samples classified as CMS3 and CMS4, all 3 MmCMS options show significant positive correlation with related human CMS subtypes, although again MmCMS-B and C classification calls display higher association to human traits compared to MmCMS-A (Supplementary Table S5).

### MmCMS-B and C (biological knowledge-based approaches) are less influenced by nonbiological factors

To assess how much non-biological factors, such as normalization methods would affect the CMS classification result of 3 options, we generated a larger collection of GEM models by combining both cohorts used in this study. APN and AP models were excluded, as the batch they were sequenced in was deeply confounded by genotype, resulting in a collection of transcriptomic data from 51 tumour samples, including 6 genotypes; A, AK, BP, BPN, KP and KPN. After batch correction using ComBat_seq, two different methods of normalisation, namely quantile and vst, were applied and thereafter CMS classification was performed using the 3 options. The results show 100% concordance between both methods for CMS calls assigned by MmCMS-B and MmCMS-C, however in line with limitations of gene-level classifiers, concordance with the gene-level MmCMS-A classifier was reduced to 92% (Supplementary Fig. 4). This suggests broad biological knowledge-based approaches based on overall gene ranking across biological pathways, rather than individual genes, are more robust and less likely to be influence by non-biological factors (12).

### Mouse CMS subtype specific biomarkers could not classify human samples

Our study suggests that individual gene-level classifiers derived from human CRC tumours perform poorly when applied to data derived from mouse CRC tumours, therefore we next assessed if CMS-specific significant genes from mouse tumours classified using our MmCMS-C method can distinguish human CMS subtypes. To this end we performed differential gene expression analysis to identify the 20 most significant CMS-specific genes for each subtype assigned using MmCMS-C (Fig. 5a). These genes were then applied to the human TCGA cohort according to the CMS subtypes, which revealed that while genes most associated with CMS4 could identify this subtype regardless of species, genes associated with mouse CMS1-3 displayed inconsistent subtype associations (Fig. 5b).

**Fig. 5:**
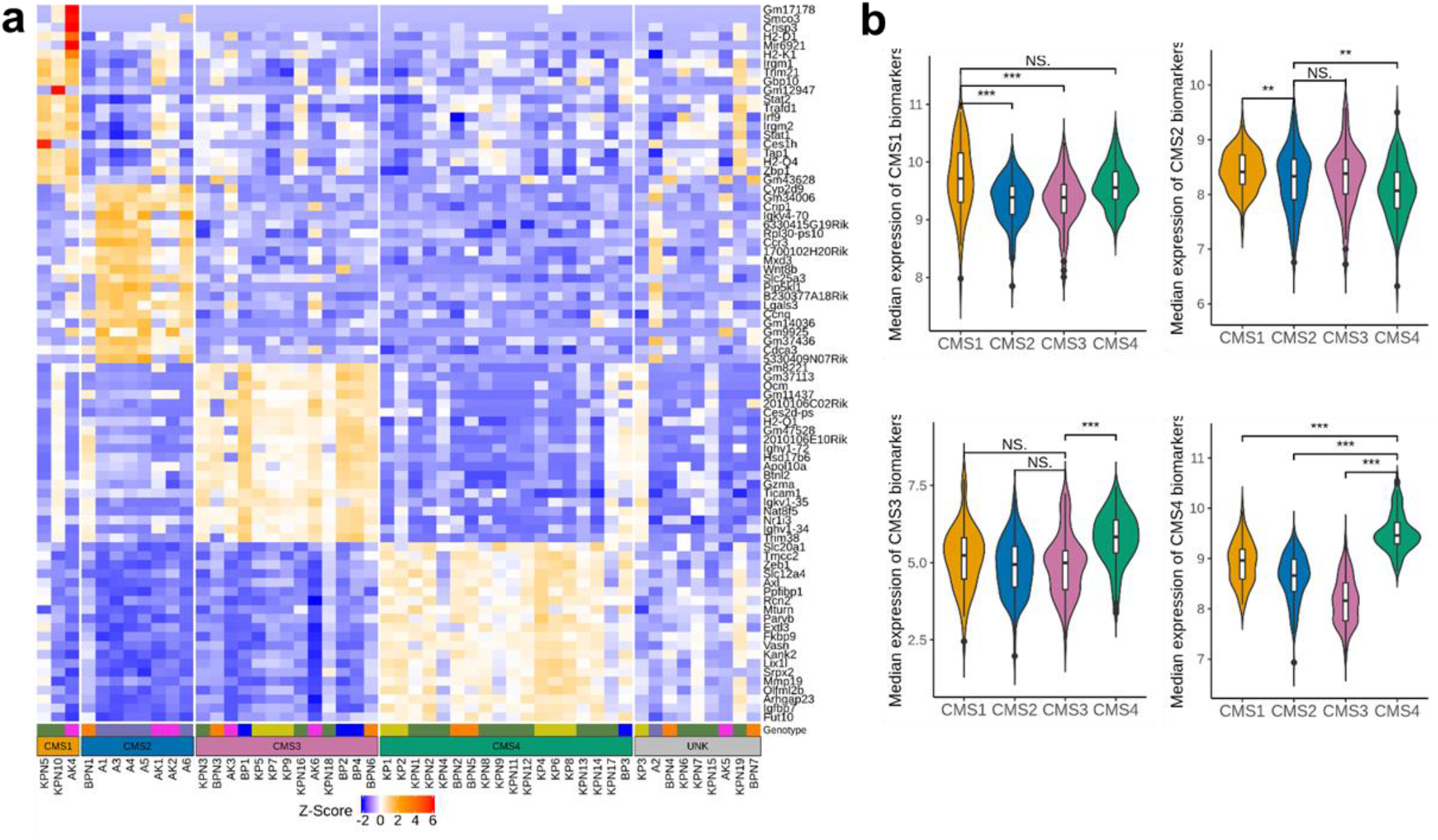
Mouse CMS subtype specific biomarkers are inconsistent across mouse and human tumour samples. **a** Heatmap of the top 20 significant differentially expressed genes (DEGs) in each CMS subtype called by MmCMS-C (Student’s t-tests in Partek Flow applied on each CMS subtype versus all other subtypes to identify DEGs). **b** Violin plot shows the median expression of the top 20 genes in human CRC samples according to CMS subtypes; DEGs (top 20) from mouse were converted to human orthologues using biomaRt. *** p ≤ 0.001; ** p ≤ 0.01; * p ≤ 0.05

## Discussion

Genetically engineered mouse models represent a valuable tool to test novel treatments that may benefit specific subtypes of tumours, making it essential to ensure the chosen models accurately recapitulate biological signalling and phenotypes underpinning human subtypes (19). Therefore, accurate and robust classification of mouse CRCs according to human subtypes is a critical step to improve disease-positioning of models and translation of findings from the pre-clinical setting. Integrity and robustness in positioning models with human cancer subtypes is critical in the era of stratified medicine, where therapeutic approaches are designed for the biology underpinning specific tumour subtypes. In order to successfully translate pre-clinical efficacies into clinical benefit, testing of therapeutics must be performed in models that are representative of specific patient subtypes. Despite its importance, dual-species classification has been limited by the lack of a reliable and standardised approach, limiting researchers’ ability to ensure faithful alignment between human tumours and preclinical models. Therefore, to address this, we developed a series of dual-species CMS classification models, named MmCMS, and an accompanying R package, which allows users to rapidly perform CMS classification of mouse tissue using three different options (A-C) of increasing complexity, from gene-level to biological pathways. To ensure that these new classifier options benefit the field, we developed a publicly available R package for MmCMS, which can be downloaded from https://github.com/MolecularPathologyLab/MmCMS. Although we have focussed on CMS in this study, data presented here provide an ideal template for the development and testing of other dual-species classifications, for subtypes such as CRC intrinsic subtypes (CRIS) (5), Braf mutant subtypes (BM) (20) and many others.

Our gene-level classifier, MmCMS-A, converts the human CMS template, embedded in CMScaller R package, to mouse orthologs and then use the NTP algorithm to carry out mouse CMS classification. The CMScaller package has been developed to enable exploration of the CMS subtypes in human pre-clinical models, particularly in cell lines, organoids and PDX tumours, to overcome the limitation of CMSclassifier’s strong dependence on gene expression derived from the tumour microenvironment (1,6). As this approach is based on individual genes, any genes lost during the process of obtaining mouse orthologues (21,22) can affect classification performance, resulting in a higher number of inaccurate or unknown calls, compared to biological knowledge-based approaches. In addition to biological differences between mouse and human, the representation and coverage of individual genes required for robust CMS classification may not be equivalent across different transcriptome profiling platforms (11), which again can lead to poor classifier performance. Recent studies have shown that classifiers based on biological pathways, rather than individual genes, have the potential to provide a more robust classification, as by using hundreds of co-ordinately expressed genes they become far less sensitive to bias that is associated with missing individual genes (9,10). This is based on the understanding that ontology/pathway-level approaches for transcriptional analyses have the advantage of identifying biologically meaningful information associated with a particular subgroup, rather than individual genes which can be confounded by issues such as intratumoural heterogeneity or technical variations associated with molecular profiling (9,11,12). In our MmCMS R package, MmCMS-B and C were developed to overcome the limitations of individual gene-based approaches and are based on ssGSEA scores from broad biological knowledge-based approaches, less influenced by nonbiological factors such as normalization methods. Correlation analysis between each CMS-related pathway in human samples shows that MmCMS-B and C are more similar to human CMS classification, using the original RF classifier, and have higher discriminatory power and classification rates, particularly for CMS2 and CMS3. Our results suggest the presence of intra-genotype CMS subtype heterogeneity, indicating that the same mutations driver events can result in variable downstream transcriptional signalling, emphasising that faithful mouse model alignment with human tumour signalling should not be based on mutation alone.

Coupled with advances in our understanding of the biology underpinning tumour development and progression, the versatility and accessibility of transcriptional signatures has seen them become a fundamental tool in the alignment of clinical phenotypes and biological signalling across human tumours and preclinical models. As therapeutics are being tested in a variety of mouse-based in vivo models, it is now even more important to ensure faithful alignment between models and human tumours and that the models we use represent the same biology during forward and reverse translation studies. Our study provides an important standardised approach for researchers to enable more reproducible and comparable classification of CRC mouse models, aligned to the biology underpinning human CRC subtypes. The identification of mouse tumours that truly mimic each human CRC subtype is essential for the proper interpretation of results, and their translation into effective human clinical trials.

## Data Availability

The dataset of 18 GEM models is available via Jackstadt et al. article (13) at ArrayExpress: E-MTAB-6363. The dataset of 39 GEM models is available from the corresponding author on reasonable request.

---

**Supplementary Fig. 1:**
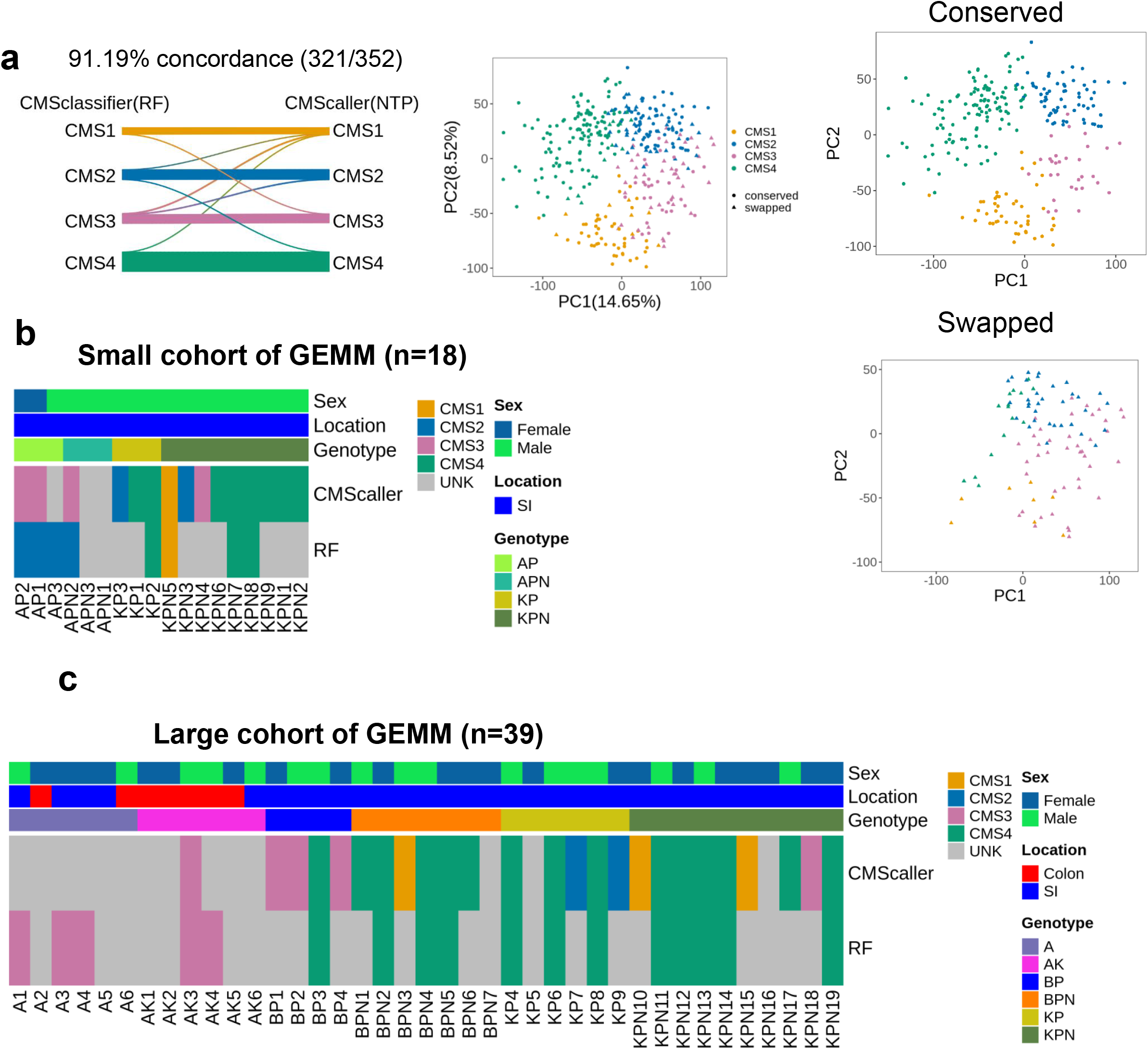
NTP method performs better than RF in CMS classification of mouse tissues. **a** Alluvial plot (top left) shows concordance between the CMS classification results of CMSclassifier (RF method) and CMScaller (NTP method) on human TCGA dataset (n=577). PCA plot depicting the clustering of TCGA data based on their CMS subtypes assigned by the RF method (top right). The PCA plot (bottom left) shows samples with the same CMS subtype called by either RF or CMScaller. The PCA plot (bottom right) shows conflicting (swapped) calls between RF and CMScaller. **b** Comparing the performance of RF and NTP methods on a small cohort of GEMMs. **c** Performance of RF and NTP methods on a large independent cohort of GEMMs. Grey colour indicates unclassified samples.

**Supplementary Fig. 2:**
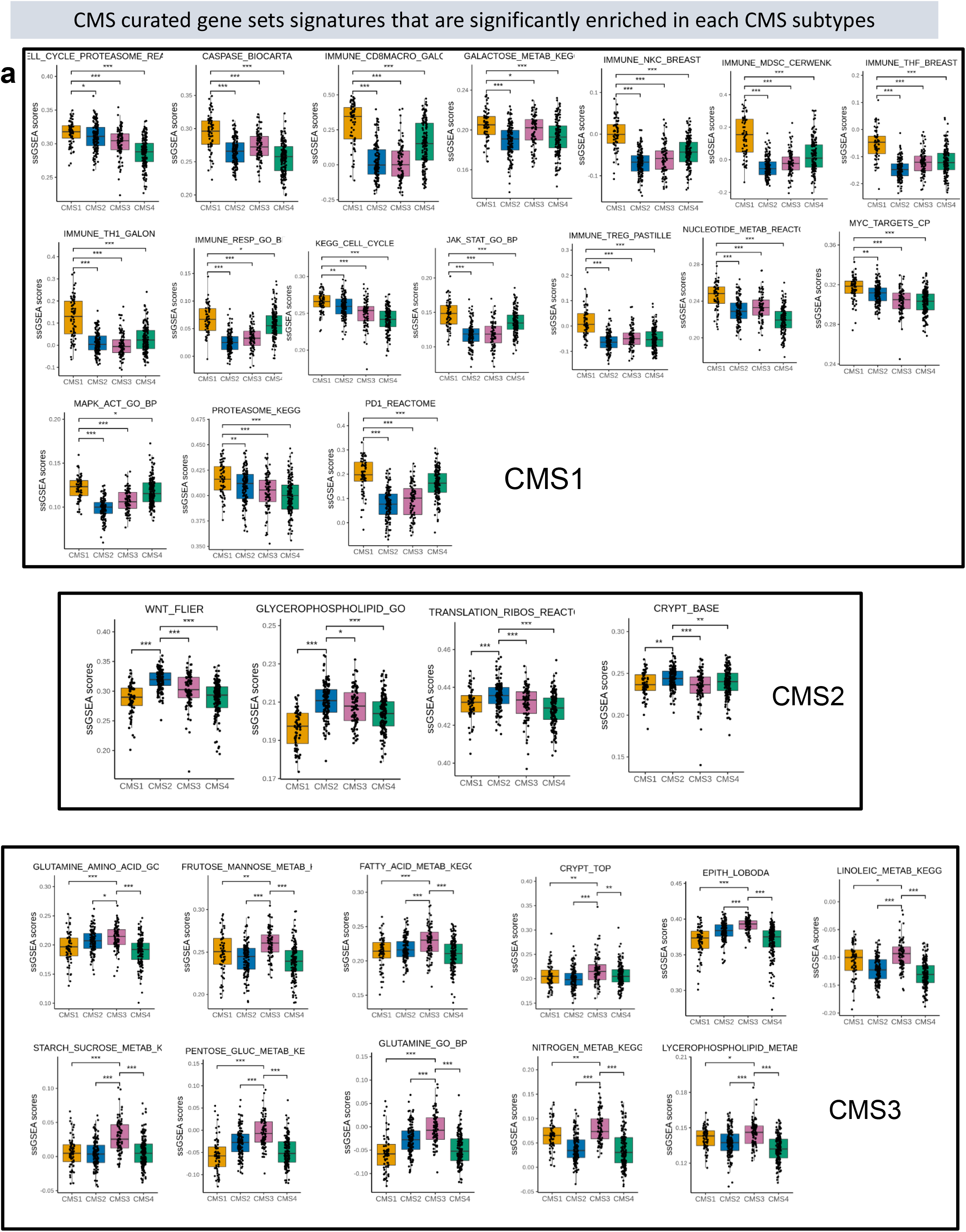
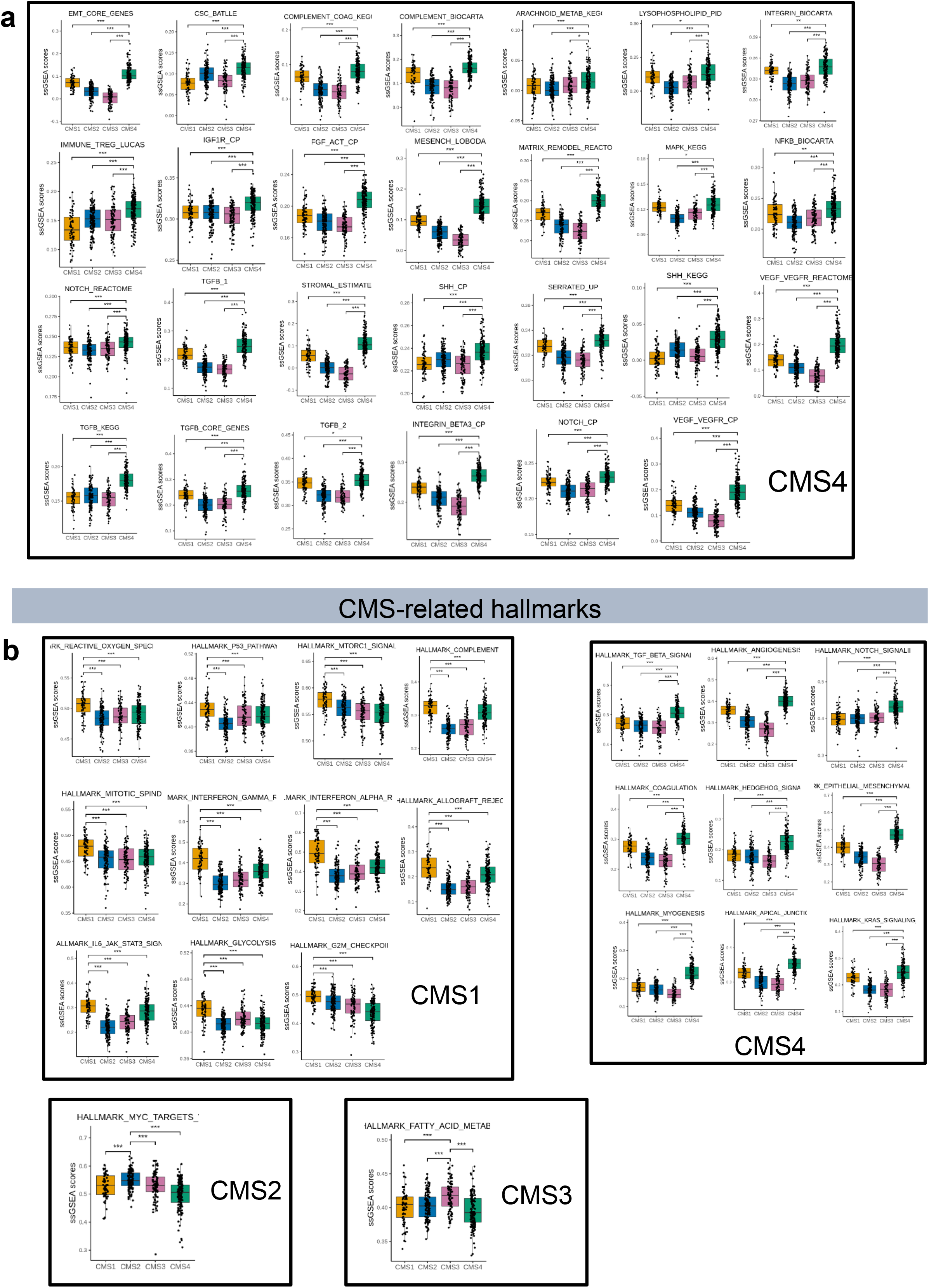
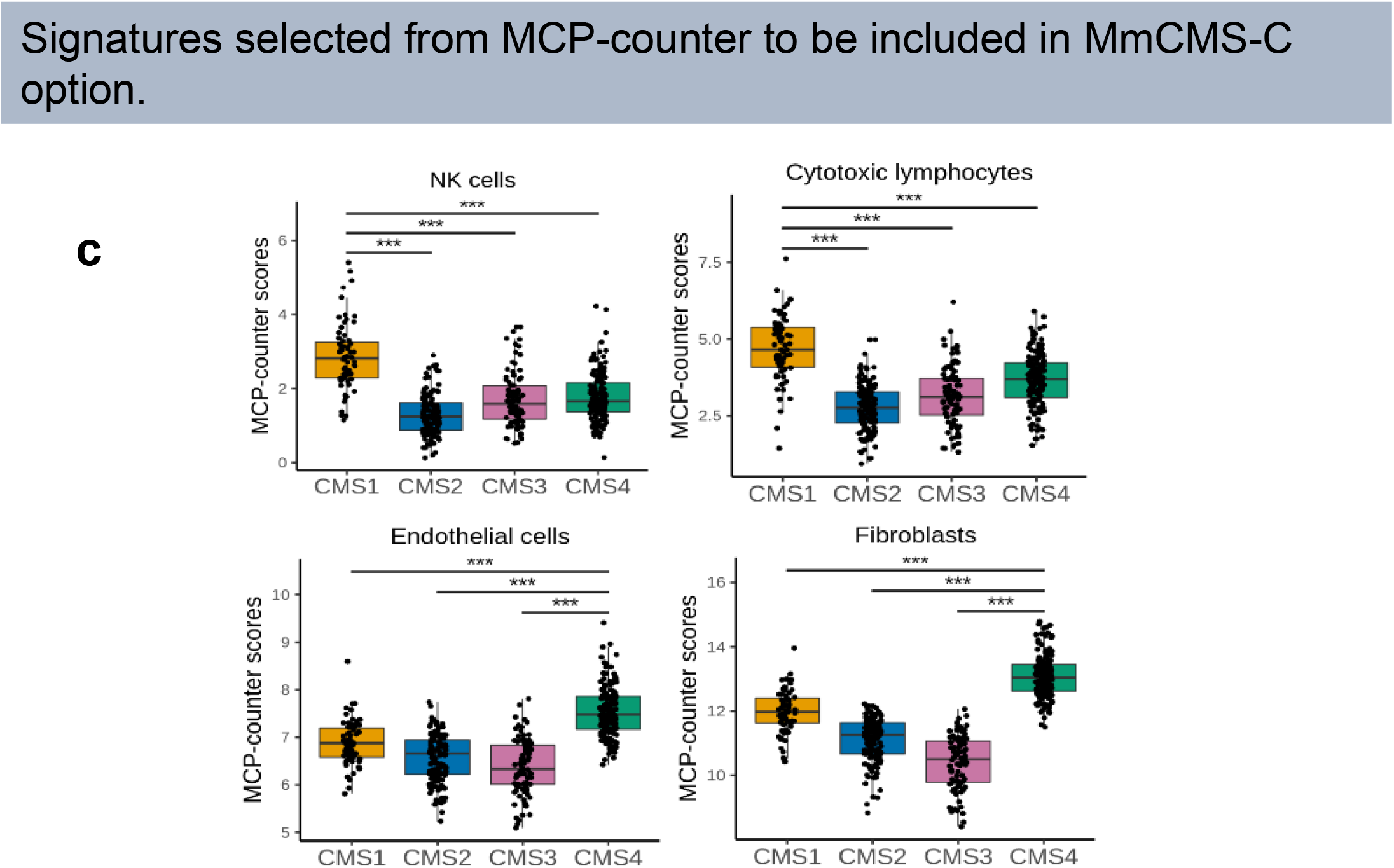
Signatures that are significantly enriched in each CMS subtypes. **a** CMS curated gene sets signatures that are significantly enriched in each CMS subtype (CMS1 samples=67, CMS2=135, CMS3=88, CMS4=155). **b** CMS-related hallmarks that are significantly enriched in each CMS subtype. **c** Four signatures from MCP-counter package were chosen to be included in the MmCMS-C option. Student *t*.*test* method embedded in the *geom_signif()* function of *ggsignif* package (v0.6.3) was used to do statistical analysis in boxplots. Horizontal line represents median values, boxes indicate the inter-quartile range and bars denote the maximum and minimum values. *** p ≤ 0.001; ** p ≤ 0.01; * p ≤ 0.05

**Supplementary Fig. 3:**
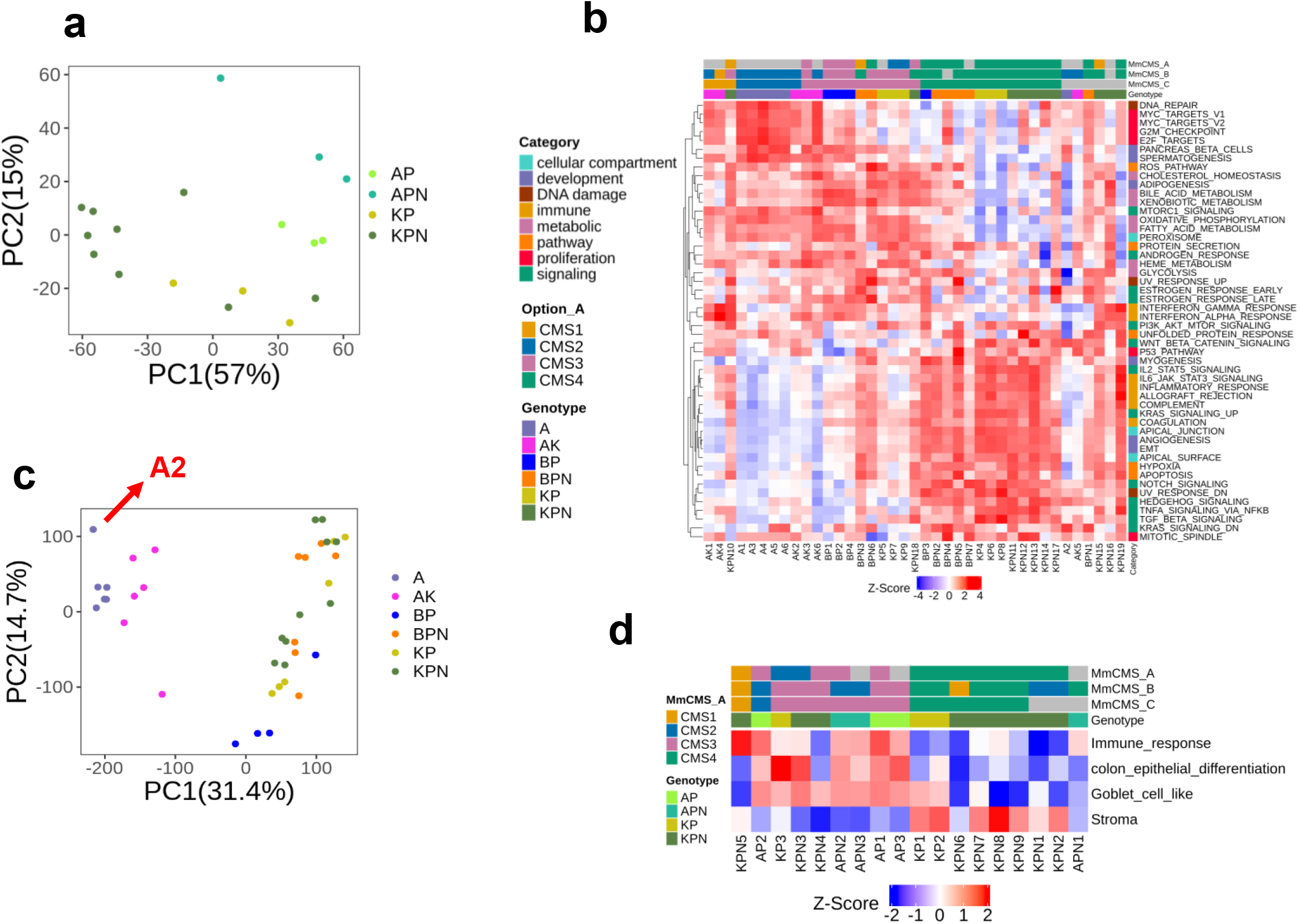
Molecular characterization of GEM models. **a** Principal component analysis (PCA) plot depicting the clustering of mouse models based on their genotype in the small cohort (Table 1). **b** heatmap of Hallmark ssGSEA score across samples in the small cohort of GEMMs (scores are z-score scaled). **c** PCA plot depicting the clustering of mouse models based on their genotype in the large GEMM cohort. **d** ssGSEA scores heatmap of immune response, colon epithelial differentiation, goblet cell–like, and stroma related gene sets across GEMMs that are aligned by genotype and CMS classification result of 3 options in the small GEMM cohort (n=18).

**Supplementary Fig. 4:**
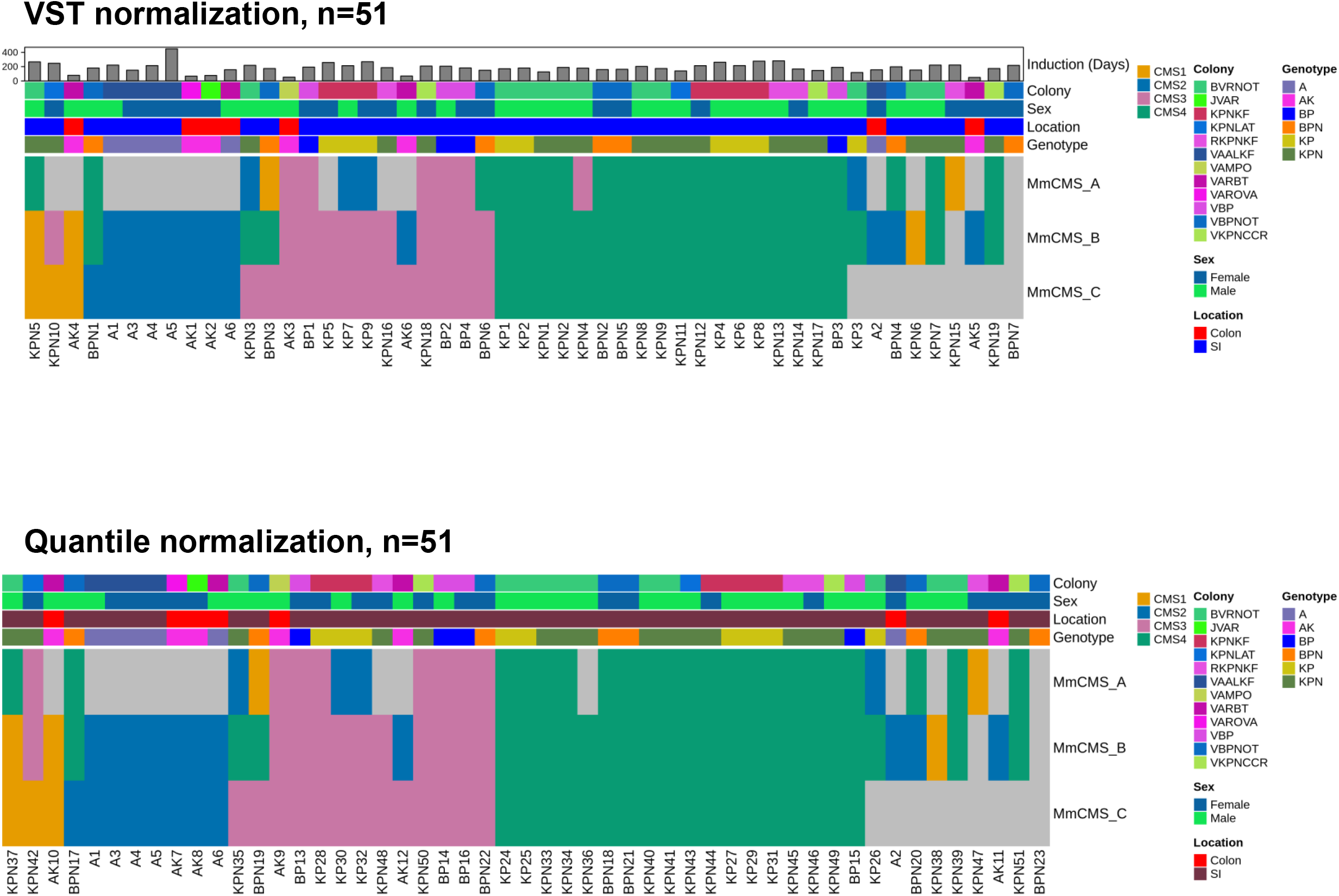
CMS classification results from the combined GEMM cohort (n=51) using the 3 classification options applied to the mouse data after normalisation by two different methods; vst in DESeq2 and quantile normalisation.

**Supplementary Table S1.**
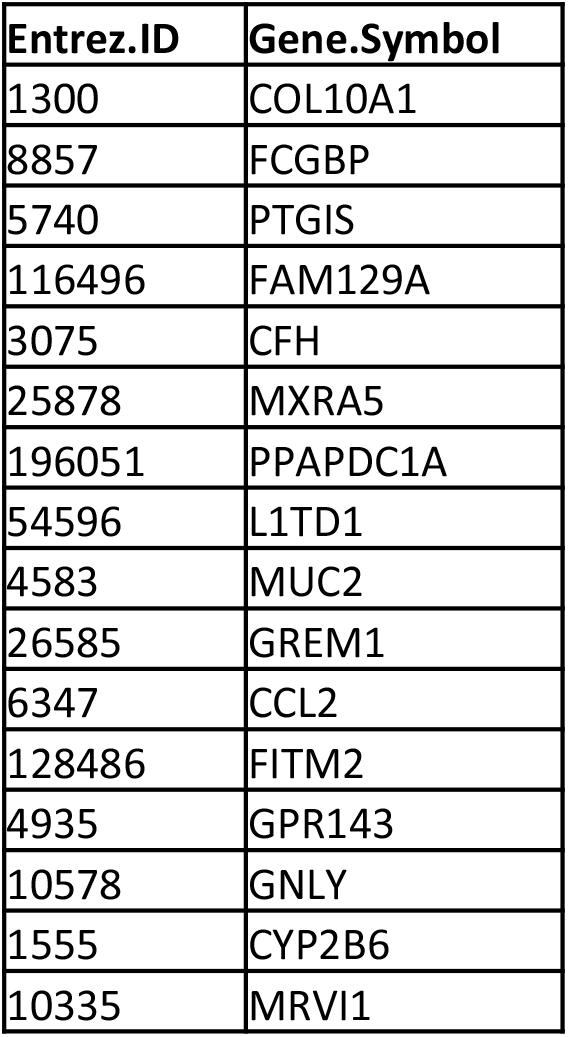
16 genes out of 273 genes in RF CMSclassifier which missed during conversion the mouse matrix to human entrez ids

| Entrez.ID | Gene.Symbol |
| --- | --- |
| 1300 | COL10A1 |
| 8857 | FCGBP |
| 5740 | PTGIS |
| 116496 | FAM129A |
| 3075 | CFH |
| 25878 | MXRA5 |
| 196051 | PPAPDC1A |
| 54596 | L1TD1 |
| 4583 | MUC2 |
| 26585 | GREM1 |
| 6347 | CCL2 |
| 128486 | FITM2 |
| 4935 | GPR143 |
| 10578 | GNLY |
| 1555 | CYP2B6 |
| 10335 | MRVI1 |

**Supplementary Table S2.** 26 genes are missing during converting human CMS to mouse orthologs due to lack of recognised orthologues

26 genes misalignment
|  |  |  |  |  |  |
| --- | --- | --- | --- | --- | --- |
| **CMS1** | CXCL5 | **CMS2** | CADPS | **CMS3** | SMAD9 |
| **CMS1** | CXCL8 | **CMS2** | COL9A3 | **CMS4** | ADAM19 |
| **CMS1** | FDXR | **CMS2** | CTSV | **CMS4** | CST1 |
| **CMS1** | IFI30 | **CMS2** | CYP2B6 | **CMS4** | CTSL |
| **CMS1** | IFI6 | **CMS2** | FER1L4 | **CMS4** | CYP2B6 |
| **CMS1** | MT1X | **CMS2** | MEP1A | **CMS4** | IFIT1 |
| **CMS1** | MT2A | **CMS3** | ANXA13 | **CMS4** | RTL8C |
| **CMS1** | PI3 | **CMS3** | CADPS | **CMS4** | SLC2A3 |
| **CMS1** | SAMD9 | **CMS3** | MUC2 |  |  |
| **CMS1** | ZBED2 | **CMS3** | SERPINA1 |  |  |

**Supplementary Table S3.** CMS-related immune genes selected using the Least Absolute Shrinkage Selector Operator (LASSO) regression model among 557 immune genes obtained from NanoString in human dataset.

| Class | Human | Mouse |
| --- | --- | --- |
| CMS1 | BCL6 | Bcl6 |
| CMS1 | CLEC5A | Clec5a |
| CMS1 | CXCR4 | Cxcr4 |
| CMS1 | DPP4 | Dpp4 |
| CMS1 | JAK2 | Jak2 |
| CMS1 | KIR3DL1 | Kir3dl1 |
| CMS1 | KIR3DL1 | Kir3dl2 |
| CMS1 | MAPK11 | Mapk11 |
| CMS1 | MBP | Mbp |
| CMS1 | PLA2G2A | Pla2g2a |
| CMS2 | BID | Bid |
| CMS2 | CD3EAP | Cd3eap |
| CMS2 | CXCL2 | Cxcl1 |
| CMS2 | IL12A | Il12a |
| CMS2 | MAPK14 | Mapk14 |
| CMS2 | NOD2 | Nod2 |
| CMS2 | TCF7 | Tcf7 |
| CMS2 | TRAF5 | Traf5 |
| CMS3 | BCL3 | Bcl3 |
| CMS3 | GFI1 | Gfi1 |
| CMS3 | IL1A | Il1a |
| CMS3 | IL22 | Il22 |
| CMS3 | IL23A | Il23a |
| CMS3 | ITLN1 | Itln1 |
| CMS3 | NOS2 | Nos2 |
| CMS3 | PTGER4 | Ptger4 |
| CMS3 | TNFRSF11A | Tnfrsf11a |
| CMS4 | BST1 | Bst1 |
| CMS4 | C1S | C1s |
| CMS4 | C7 | C7 |
| CMS4 | CD81 | Cd81 |
| CMS4 | ENTPD1 | Entpd1 |
| CMS4 | FCGRT | Fcgrt |
| CMS4 | FN1 | Fn1 |
| CMS4 | MARCO | Marco |
| CMS4 | PDGFRB | Pdgfrb |
| CMS4 | STAT5B | Stat5b |
| CMS4 | THY1 | Thy1 |
| CMS4 | ZEB1 | Zeb1 |

**Supplementary Table S4.** The number of samples per CMS subtypes assigned by MmCMS-C in small cohort.

| Option_C approach |  |  |  |  |  |
| --- | --- | --- | --- | --- | --- |
|  | CMS1 | CMS2 | CMS3 | CMS4 | NA |
| KPN | 1 | - | 2 | 4 | 2 |
| KP | - | - | 1 | 2 | - |
| APN | - | - | 2 | - | 1 |
| AP | - | 1 | 2 | - | - |

**Supplementary Table S5.** Correlation analysis between human CRC and each mouse option in Figure 4 for each CMS subtype

| Small cohort (n=18) |  |  |  |
| --- | --- | --- | --- |
| Human | Mouse | estimate | p.value |
| RF_CMS1 | A_CMS1 | 0.696878 | 3.79E-05 |
| RF_CMS1 | B_CMS1 | 0.557433 | 0.002058 |
| RF_CMS1 | C_CMS1 | 0.696878 | 3.79E-05 |
| RF_CMS2 | A_CMS2 | -0.23087 | 0.237203 |
| RF_CMS2 | B_CMS2 | 0.792418 | 5.02E-07 |
| RF_CMS2 | C_CMS2 | 0.813795 | 1.39E-07 |
| RF_CMS3 | A_CMS3 | 0.840478 | 2.19E-08 |
| RF_CMS3 | B_CMS3 | 0.914454 | 1.03E-11 |
| RF_CMS3 | C_CMS3 | 0.905587 | 3.51E-11 |
| RF_CMS4 | A_CMS4 | 0.807208 | 2.10E-07 |
| RF_CMS4 | B_CMS4 | 0.922386 | 3.03E-12 |
| RF_CMS4 | C_CMS4 | 0.890457 | 2.22E-10 |

| Large cohort (n=39) |  |  |  |
| --- | --- | --- | --- |
| Human | Mouse | estimate | p.value |
| RF_CMS1 | A_CMS1 | -0.15213 | 0.439622 |
| RF_CMS1 | B_CMS1 | 0.36814 | 0.053916 |
| RF_CMS1 | C_CMS1 | 0.2099 | 0.283699 |
| RF_CMS2 | A_CMS2 | 0.160997 | 0.413108 |
| RF_CMS2 | B_CMS2 | 0.673605 | 8.53E-05 |
| RF_CMS2 | C_CMS2 | 0.706416 | 2.66E-05 |
| RF_CMS3 | A_CMS3 | 0.906198 | 3.24E-11 |
| RF_CMS3 | B_CMS3 | 0.864359 | 3.07E-09 |
| RF_CMS3 | C_CMS3 | 0.909069 | 2.20E-11 |
| RF_CMS4 | A_CMS4 | 0.898072 | 9.10E-11 |
| RF_CMS4 | B_CMS4 | 0.922993 | 2.75E-12 |
| RF_CMS4 | C_CMS4 | 0.927934 | 1.19E-12 |

